# CHI3L1 enhances melanoma lung metastasis via regulation of t cell co-stimulators and CTLA-4/B7 axis

**DOI:** 10.1101/2022.06.23.497390

**Authors:** Bing Ma, Suchitra Kamle, Bedia Akosman, Hina Khan, Chang Min Lee, Chun Geun Lee, Jack A. Elias

**Affiliations:** Molecular Microbiology and Immunology, Brown University, 185 Meeting St., Providence, Rhode Island; Divsion of Hematology-Oncology, Warren Alpert Medical School, Brown University, Providence, Rhode Island; Department of Medicine, Brown University, Providence, Rhode Island

**Keywords:** Chitinase 3-like-1, melanoma, metastasis, ICOS, ICOSL, CTLA-4, B7-1, B7-2, CD28, anti-CHI3LI, bispecific antibodies

## Abstract

ICOS/ICOSL and CD28/B7-1/B7-2 are T cell co-stimulators and CTLA-4 is an immune checkpoint inhibitor that play critical roles in the pathogenesis of neoplasia. Chitinase 3-like-1 (CHI3L1) is induced in many cancers where it portends a poor prognosis and contributes to tumor metastasis. Here we demonstrate that CHI3L1 inhibits the expression of ICOS, ICOSL and CD28 while stimulating CTLA-4 and the B7 moieties in melanoma lung metastasis. We also demonstrate that RIG-like helicase innate immune activation augments T cell co-stimulation, inhibits CTLA-4 and suppresses pulmonary metastasis. At least additive antitumor responses were seen in melanoma lung metastasis treated with anti-CTLA-4 and anti-CHI3L1 antibodies in combination. Synergistic cytotoxic T cell-induced tumor cell death and the heightened induction of the tumor suppressor PTEN were seen in co-cultures of T and tumor cells treated with bispecific antibodies that target both CHI3L1 and CTLA-4. Thus, CHI3L1 contributes to pulmonary metastasis by inhibiting T cell co-stimulation and stimulating CTLA-4. The simultaneous targeting of CHI3L1 and the CTLA-4 axis with individual and, more powerfully with bispecific antibodies, represent promising therapeutic strategies for pulmonary metastasis.

## Introduction

Neoantigens are generated during the development and progression of many cancers (1). In the majority of circumstances, the new antigens are recognized as “non-self” allowing the host to mount T cell-mediated “non self” immune responses that can control and or kill tumor cells (1). The T cell activation that occurs during these responses requires 2 signals. The first (signal 1), is provided by the interaction of the antigen/peptide with the T cell receptor (TCR) and major histocompatibility complex II (MHC II) on the surface of the antigen presenting cell (APC). The second is an antigen-independent co-stimulation which is mediated by moieties such as ICOS and ICOS ligand (ICOSL) and CD28 and its ligands B7-1 and B7-2 (also called CD80 and CD86). T cell activation is also highly regulated by immune checkpoints (ICP) molecules, the receptors and their ligands such as programed death (PD)-1 and its ligands PD-L1 and PD-L2; cytotoxic T lymphocyte antigen 4 (CTLA-4) and its ligands B7-1 and B7-2; and lymphocyte activation gene 3 protein (LAG3) and its ligand HLA class II (2). These are important findings because it is now known that cancers activate ICP pathways to inhibit host anti-tumor responses (3, 4). In addition, immune checkpoint molecule blocking antibodies targeting PD-1, PD-L1, and CTLA-4 have been successfully applied as therapeutics for a number of malignancies (2, 5). However, the moieties that regulate the expression of co-stimulators and inhibitory ICP have not been adequately defined.

CTLA-4 is a homodimeric glycoprotein receptor that is related to CD28. It is expressed on T cells within 1 hour of antigen exposure (6) and inhibits T cell signaling by out-competing CD28 for binding to B7-1 and B7-2. It is also constitutively expressed on the surface of regulatory T cells (T regs) where it is required for contact-mediated immunosuppression (7). Antibodies against CTLA-4 diminish tumor growth in murine neoplastic modeling systems (8) and are effective as mono therapies in advanced melanoma, colorectal and renal cell cancers (6). However, the processes that regulate the production of CTLA-4 have not been adequately defined.

Chitinase 3-like-1 (CHI3L1; also called YKL-40), the prototypic chitinase-like protein (CLP), is expressed in various cells that include myeloid cells (macrophages, neutrophils, monocytes), endothelial and epithelial cells and is stimulated by a variety of inducers of injury, remodeling or inflammation (9). Studies from our laboratory and others have demonstrated a pleiotropic role of CHI3L1 that inhibits apoptotic cell death, stimulates alternative macrophage activation and Th2 cell differentiation and inhibits oxidative stress-induced injury. It also reduces inflammasome activation, regulates TGF-β1 elaboration, and enhances antibacterial responses mainly through MAP Kinase, Akt/PKB and Wnt/β-catenin signaling (10–14). Elevated levels of circulating and tissue expression of CHI3L1 have been reported in a wide variety of diseases characterized by inflammation, fibrosis and tissue remodeling (9, 15–18). The levels of circulating CHI3L1 are also increased in cancers of the prostate, colon, lung, rectum, ovary, kidney, breast, glioblastomas and malignant melanoma where they frequently correlate with disease progression and inversely with disease-free interval and survival (19–27). Studies from our laboratory have demonstrated that induced expression of CHI3L1 plays a critical role in the generation of a metastasis permissive tumor microenvironment (28) and the inhibition of CHI3L1 via RIG-like helicase (RLH) innate immune activation can also inhibit metastatic spread in the lung (29). Our studies also identified that CHI3L1 is a potent stimulator of the PD-1-PD-L1/2 inhibitory immune checkpoint molecules (30). However, the mechanism(s) by which CHI3L1 contributes to tumor development and progression have not been fully defined and the degree to which CHI3L1 contributes to its tumorigenic effects via activating T cell co-stimulators or the CTLA-4-B7 axis have not been investigated. Although RLH activation plays a critical role in the responsiveness to immune checkpoint blockade (30), if this responsiveness is mediated through RLH inhibition of CHI3L1 has not been addressed.

We hypothesized that CHI3L1 plays an essential role in pulmonary metastasis via regulation of T cell co-stimulation and inhibitory ICP. To test this hypothesis, we determined if CHI3L1 regulates co-stimulatory and or ICP pathways. These studies demonstrate that CHI3L1 inhibits ICOS, ICOS ligand (ICOSL) and CD28 while stimulating B7-1, B7-2 and CTLA-4. They also demonstrate that RLH innate immune activation augments co-stimulation, inhibits CTLA-4 and suppresses pulmonary metastasis. At least additive antitumor effects were seen in the mice with melanoma lung metastasis treated simultaneously with individual antibodies against CTLA-4 and CHI3L1. In addition, synergistic cytotoxic T cell (CTL)-induced tumor cell death and the induction of the tumor suppressor PTEN were seen in a T cell-tumor co-culture treated with bispecific antibodies that target CHI3L1 and CTLA-4. These studies demonstrate that CHI3L1 contributes to the initiation and or progression of metastatic tumors in the lung via the regulation of co-stimulation and CTLA-4-mediated immunosuppression. They also strongly support the notion that the simultaneous targeting of CHI3L1 and CTLA-4 with monospecific antibodies and, even more powerfully with bispecific antibodies, represent novel and attractive therapeutic strategies for lung metastasis.

## Results

### Pulmonary melanoma metastasis inhibits the Co-stimulators ICOS, ICOSL and CD28

Studies were undertaken to determine if the expression of T cell co-stimulators such as ICOS and ICOSL was altered by the melanoma lung metastasis. After C57BL/6 mice were challenged with freshly prepared B16-F10 (B16) melanoma cells or vehicle controls (PBS), the expression of ICOS, ICOSL and CD28 in the lung was assessed 2 weeks later. We noted decreased expression of ICOS and ICOSL in lungs with melanoma metastasis compared to vehicle controls (Figure 1, A and B). This regulation was not specific for the ICOS moieties because the levels of mRNA encoding CD28 were also reduced (Figure 1C). In all cases, these changes in mRNA were associated with comparable alterations in the levels of costimulatory proteins (Figure 1D). They were prominent in CD11b+ and SPC+ cells (Figure 1, E and F; Figure S1, A and B). When viewed in combination, these studies demonstrate that melanoma lung metastasis is associated with significantly suppressed expression of T cell co-stimulators including ICOS, ICOSL and CD28.

**Figure 1.**
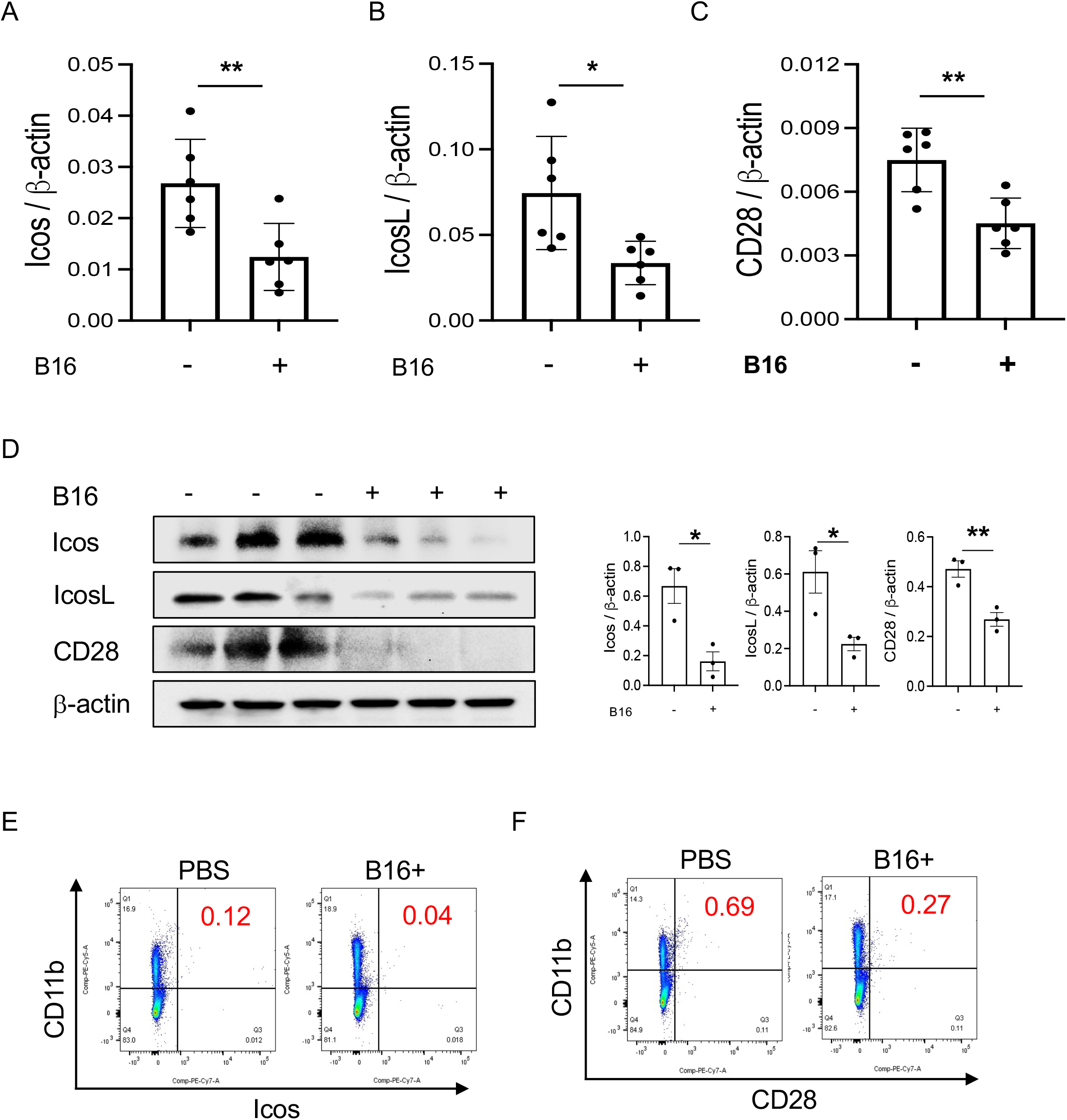
Pulmonary melanoma metastasis inhibits T cell co-stimulation. WT mice were given B16-F10 (B16) melanoma cells or control vehicle (PBS) and evaluated 2 weeks later. (AC) Real-time RT-PCR (RT-PCR) was used to quantitate the levels of mRNA encoding ICOS, ICOSL and CD28 in the lungs from mice treated intravenously with PBS (B16 -) or B16 cells (B16 +). Each dot represents an evaluation in an individual animal. (D) Western blot and densitometry evaluations of ICOS, ICOSL and CD28 accumulation in lungs from mice treated with PBS (B16 -) or B16 cells (B16 +). (E and F) FACS evaluations quantitating the accumulation of ICOS and CD28 in cell populations from lungs of mice treated with B16 cells (B16+) or vehicle control (B16 -). These evaluations used specific markers of airway myeloid cells. The values in panel A represent the mean±SEM of the noted evaluations. Panels D, E and F are representative of a minimum of 2 similar evaluations. \**P*<0.05, \*\**P*<0.01 (*t*-Test).

### Pulmonary melanoma metastasis stimulates CTLA-4 and its ligands

Studies were next undertaken to determine if the expression of inhibitory ICP, such as CTLA-4, were altered by the melanoma lung metastasis. After C57BL/6 mice were challenged with B16 melanoma cells or vehicle control, the expression of CTLA-4 was evaluated 2 weeks later. We noted a significant stimulation of CTLA-4 and its ligands B7-1 and B7-2 was noted in the lungs with melanoma metastasis (Figure 2, A-C). We also appreciated comparable alterations in the levels of CTLA-4 and its B7 ligands (Figure 2D) expression in CD11b+ and CD3+ cells and, to a lesser extent, SPC+ and CC10 + cells (Figure 2, E and F; Figure S1, C and D and Figure S2A). This regulation was not specific for the CTLA-4 because PD-1, PD-L1 and LAG6 were similarly induced as described by our laboratory (30). It was also not specific for ligands of B7-1 and or B7-2 because CD28, which also binds these moieties, was inhibited in this setting as noted above. These studies demonstrate that melanoma lung metastasis is associated with enhanced expression of CTLA-4 and B7-1 and B7-1.

**Figure 2.**
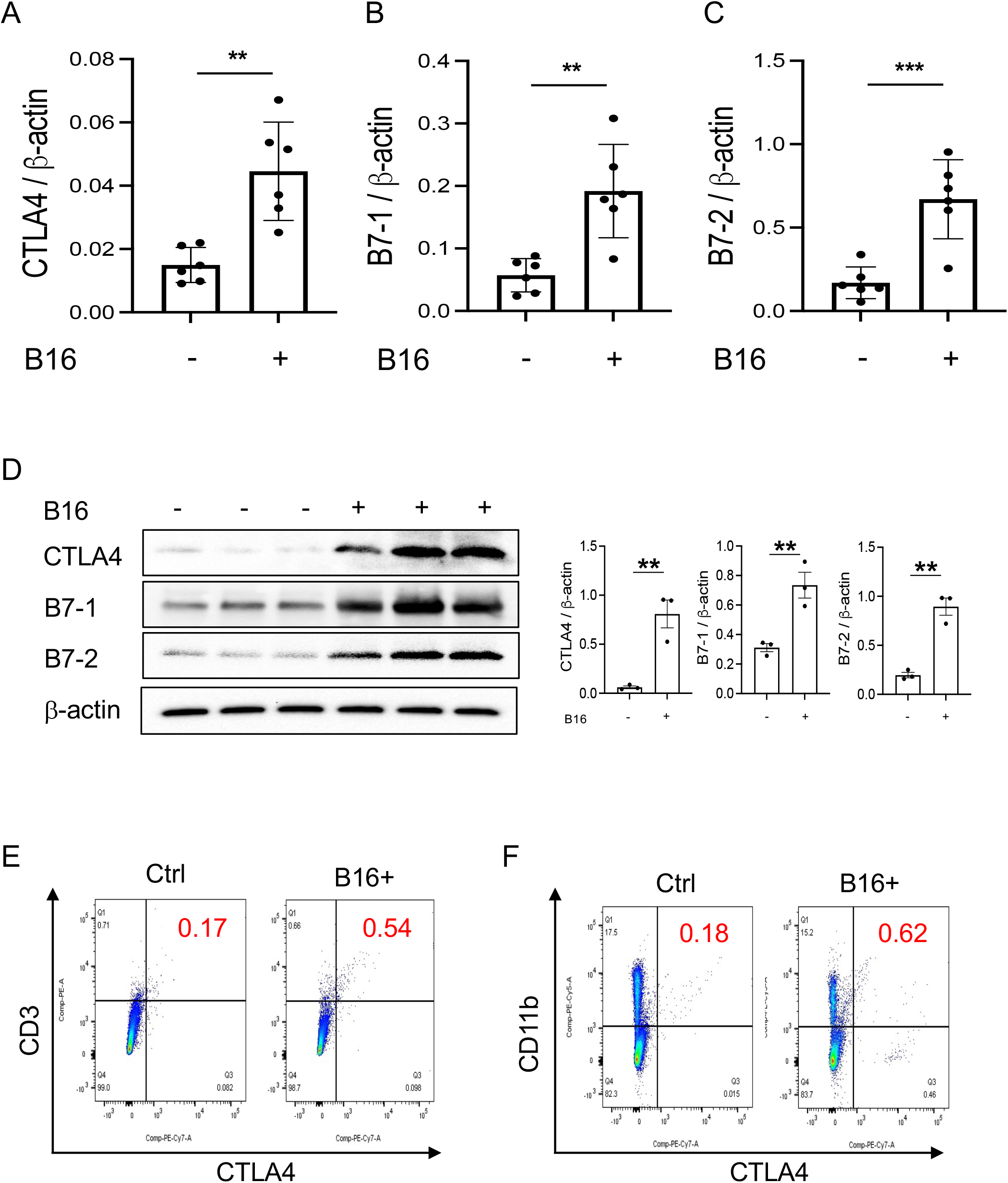
Chi3l1 plays a critical role in B16 melanoma stimulation of pulmonary CTLA-4 and its ligands. WT mice were given B16-F10 (B16) melanoma cells or control vehicle (PBS) and evaluated 2 weeks later. (A-C) RT-PCR was used to quantitate the levels of mRNA encoding CTLA-4, B7-1 and B72 in the lungs from mice treated intravenously with PBS vehicle (B16 -) or B16 cells (B16 +). Each dot represents an evaluation in an individual animal. (D) Western blot and densitometry evaluations of CTLA-4, B7-1 and B7-2 accumulation in lungs from WT mice treated with vehicle (B16-) or B16 cells (B16 +). (E and F) FACS analysis demonstrating the induction of CTLA-4 by CD11b+ and CD3+ cells. The plotted values in panels A-C represent the mean±SEM of the noted evaluations represented by the individual dots. Panels D and E are representative of a minimum of 2 similar evaluations. \*\**P*<0.01, \*\*\**P*<0.001 (*t*-Test).

### CHI3L1 plays an important role in in melanoma metastasis inhibition of co-stimulation and the induction of CTLA-4 and its ligands

To begin to determine the role(s) of CHI3L1 in the inhibition of co-stimulation and induction of CTLA-4 and its ligands, we compared the levels of ICOS, ICOSL, CD28, CTLA-4, B7-1 and B7-2 in wild type (WT) and CHI3L1 (*Chil1*) null mutant (*Chil1*^-/-^) mice that were randomized to receive B16 cells or vehicle control. As can be seen in Figure 3 in panels A and B, the inhibition of ICOS, ICOSL and CD28 and the stimulation of CTLA-4 and its ligands were abrogated in mice with null mutations of CHI3L1 (*Chil1*^-/-^). In accord with these findings, monoclonal anti-CHI3L1 antibody (called FRG antibody) treatment significantly rescued the B16 cell inhibition of ICOS, ICOSL and CD28 and stimulation of CTLA-4, B7-1 and B7-2 (Figure 3, C and D). In addition, the ability of B16-F10 cells to stimulate CTLA-4, B7-1 and B7-2 were further exaggerated in lungs with transgenic CHI3L1 overexpression (Figure 3E). When viewed in combination, these studies demonstrated an essential role of CHI3L1 in melanoma inhibition of T cell co-stimulation and the induction of CTLA-4 and its ligands.

**Figure 3.**
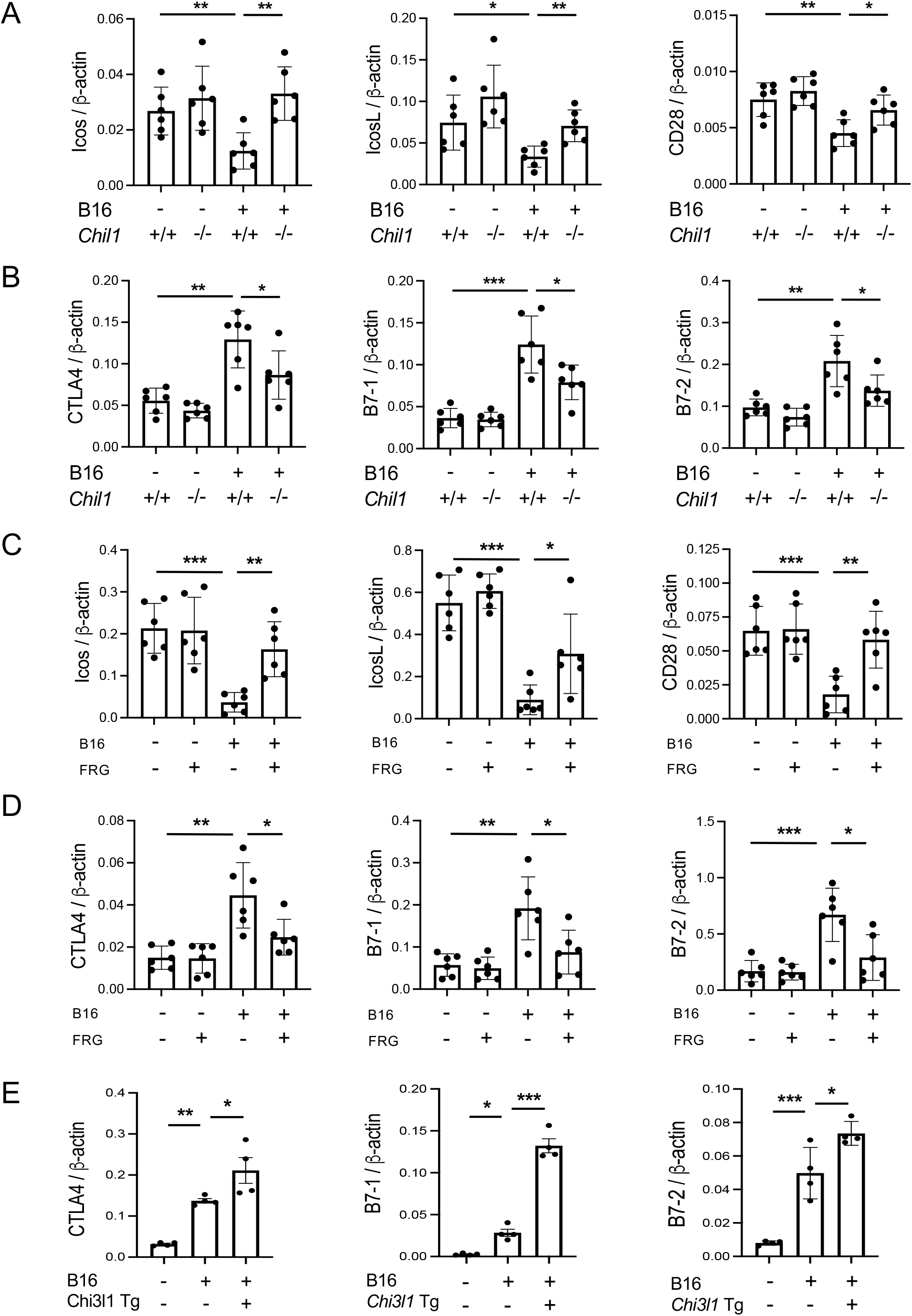
CHI3L1 plays a critical role in B16 cell regulation of T cell co-stimulators and the induction of CTLA-4 and its ligands. WT (*Chil1*^+/+^) and Chi3l1 null (*Chil1*^-/-^) mice were used to evaluate the expression and accumulation of co-stimulators (ICOS, ICOS L CD28) and CTLA-4, B7-1 and B7-2 in the lung. (A) RT-PCR was used to quantitate the levels of mRNA encoding ICOS, ICOS L and CD28 in the lungs from WT mice and *Chil1*^-/-^ mice randomized to receive B16 cells (B16+) or control vehicle (B16-). (B) RT-PCR was used to quantitate the levels of mRNA encoding CTLA-4, B7-1and B7-2 in the lungs from WT mice and *Chil1*^-/-^ mice randomized to receive B16 cells (B16+) or control vehicle (B16-). (C and D) RT-PCR was used to quantitate the levels of mRNA encoding ICOS, ICOS L, CD28 CTLA-4, B7-1 and B7-2 in the lungs from WT mice randomized to receive B16 cells (B16+) or vehicle control (B16-) and randomized to receive the FRG antibody (FRG+) or an isotype antibody control (FRG-). (E) RT-PCR was used to quantitate the levels of mRNA encoding CTLA-4, B7-1 and B7-2 in the lungs from WT mice and Chi3l1 transgenic (*Chi3l1* Tg) mice randomized to receive B16 cells (B16+) or vehicle control (B16-). In all panels each dot represents the evaluation in an individual animal. The values in these panels represent the mean±SEM of the noted evaluations represented by the individual dots. \**P*<0.05. \*\**P*<0.01, \*\*\**P*<0.005 (ANOVA with multiple comparisons).

### Transgenic CHI3L1 inhibits co-stimulation and stimulates CTLA-4 and its ligands

In studies noted above, melanoma lung metastasis inhibits T cell co-stimulation and stimulates ICPI such as CTLA-4 and its ligands and CHI3L1 plays an essential role in these regulatory events. Since CHI3L1 null mutation or α-CHI3L1 neutralizing antibody treatment reduced metastatic spread (28, 29), they do not determine if the inhibition of ICOS and ICOSL and or the induction of CTLA-4 are due to the direct effects of CHI3L1 or due to the decreased metastatic spread of melanoma. To test this notion, we determined if CHI3L1 inhibits co-stimulation and stimulates the CTLA-4 axis without melanoma metastasis. In these experiments, the expression and accumulation of ICOS, ICOSL CD28, CTLA-4, B7-1 and B7-2 were evaluated in lungs from WT mice and CHI3L1 overexpressing transgenic mice (Tg). These studies demonstrated that CHI3L1 itself is a potent inhibitor of ICOS, ICOSL and CD28 and a stimulator of CTLA-4 and B7-1 and B7-2 in lungs from CHI3L1 Tg mice when compared to WT controls (Figure 4, A-D). FACS evaluations and immunohistochemistry (IHC) demonstrated that transgenic CHI3L1 specifically stimulated CTLA-4 in many cells including CD3+ T cells, epithelial cells, and macrophages (Figure 4, D and E; Figure S2B). These studies demonstrate that the inhibition of ICOS, ICOSL and CD28 and stimulation of CTLA-4 and its ligands in melanoma lung metastasis is mediated, at least in part, by a tumor-independent effect of CHI3L1.

**Figure 4.**
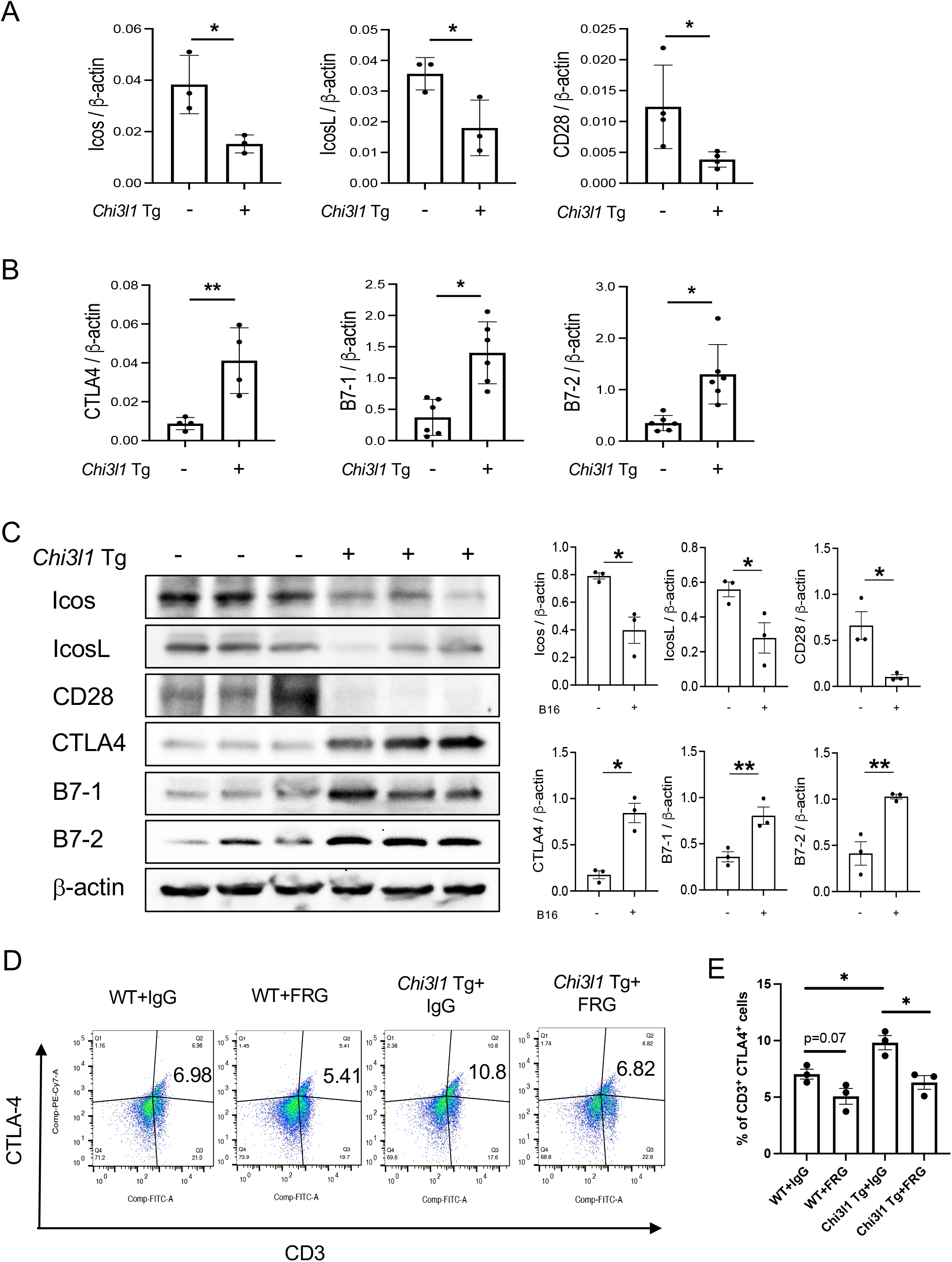
Chi3l1 inhibits co-stimulation and stimulates pulmonary CTLA-4 in the normal lung. WT and Chi3l1 Tg (+) mice were used to evaluate the effects of transgenic Chi3l1 on costimulation and the CTLA-4 axis. (A) RT-PCR was used to quantitate the levels of mRNA encoding ICOS, ICOS L and CD28 in the lungs from WT mice and Chi3l1 Tg+ mice. (B) RT-PCR was used to quantitate the levels of mRNA encoding CTLA-4, B7-1and B7-2 in the lungs from WT mice and Chi3l1 Tg+ mice. (C) Western blot and densitometry evaluations of ICOS, ICOSL, CD28 B7-1 and B7-2 accumulation in lungs from WT and Chi3l1 Tg+ mice. (D) FACS evaluations quantitating the accumulation of CTLA-4 in CD3+ cells from WT mice and Chi3l1 Tg+ mice. (E) Cumulative FACS evaluations quantitating the accumulation of CTLA-4 in CD3+ cells from WT and Chi3l1 Tg+ mice. The plotted values in panels A, B and E represent the mean ± SEM of the noted evaluations represented by the individual dots. Panel C is representative of a minimum of 2 similar evaluations. \**P*<0.05, \*\**P*<0.01 (*t*-Test for panels A and B, ANOVA with multiple comparisons for panels E).

### Rig-like helicase (RLH) activation augments costimulation and inhibits CTLA-4, B7-1 and B7-2

In previous studies, we demonstrated that activation of retinoic acid inducible gene I (RIG-I) and RIG-like helicase (RLH) pathway with Poly(I:C) treatment impressively inhibits CHI3L1 expression and pulmonary melanoma metastasis (29). RIG-I activation and mitochondrial antiviral signaling molecule (MAVS) are also reported to play an important role in many immune checkpoint inhibitor blockade-induced anti-tumor responses (31). Thus, we determined if RLH activation with Poly(I:C) treatment alters the ability of CHI3L1 to inhibit ICOS, ICOSL and CD28 and stimulates CTLA-4 and its ligands. Poly(I:C) ameliorated melanoma-inhibition of ICOS, ICOSL and CD28 while abrogating the ability of B16 cells to stimulate CTLA-4 (Figure 5, A and B). Flow cytometric evaluation also showed that T cell expression of CTLA-4 was prominently reduced while macrophage expression of CD28 was augmented by poly(I:C) treatment (Figure 5, C and D). The stimulation of ICOS, ICOSL, and CD28 and the inhibition of CTLA-4 and B7-1 and B7-2 by Poly(I:C) were dependent, at least in part, on CHI3L1 because the transgenic overexpression of CHI3L1 that is not regulated by the RLH activation significantly ameliorated Poly(I:C)-induced inhibition of the co-stimulators and inhibitory ICP stimulation (Figure 6, A - F). These studies highlight the ability of RLH activation that inhibits CHI3L1 expression and, in turn, eliminate the ability of CHI3L1 to inhibit co-stimulation and augment the expression and accumulation of inhibitory ICP molecules as shown in previous studies (30).

**Figure 5.**
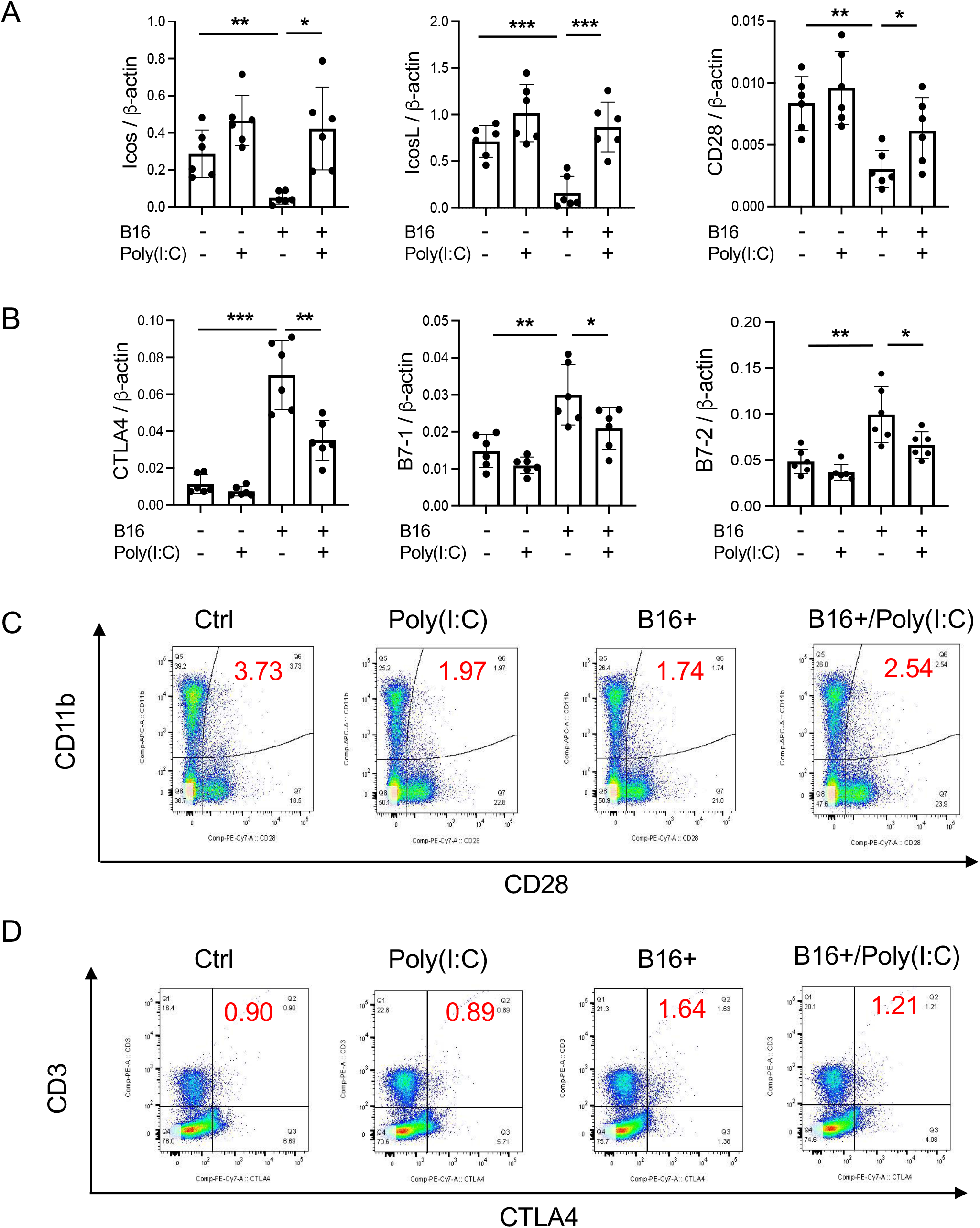
Rig-like helicase (RLH) activation augments co-stimulation and inhibits CTLA-4, B7-1 and B7-2. WT mice were used to evaluate the effects of Rig-like helicase activation with Poly (I:C) on co-stimulation and the CTLA-4 axis. (A) RT-PCR was used to quantitate the levels of mRNA encoding ICOS, ICOS L and CD28 in the lungs from WT mice that received B16 cells (B16+) or vehicle control (B16-) and were treated with Poly (I:C) (+) or its vehicle control (Poly(I:C)-). (B) RT-PCR was used to quantitate the levels of mRNA encoding CTLA-4, B7-1 and B7-2 in the lungs from WT mice that received B16 cells (B16+) or vehicle control (B16-) and were treated with Poly (I:C) (+) or its vehicle control (Poly(I:C)-). (C) FACS evaluations comparing the expression of CTLA-4 on CD3+ cells from WT mice that received B16 cells (B16+) or vehicle control (B16-) and were treated with Poly (I:C) or its vehicle control. (D) FACS evaluations comparing the expression of CD28 on CD11b+ cells from WT mice that received B16 cells (B16+) or vehicle control (B16-) and were treated with Poly (I:C) or its vehicle control. The plotted values in panels A and B represent the mean ± SEM of the noted evaluations represented by the individual dots. Panels C and D are representative of a minimum of 2 similar evaluations. \**P*<0.05, \*\**P*<0.01, \*\*\**P*<0.001 (ANOVA with multiple comparisons).

**Figure 6.**
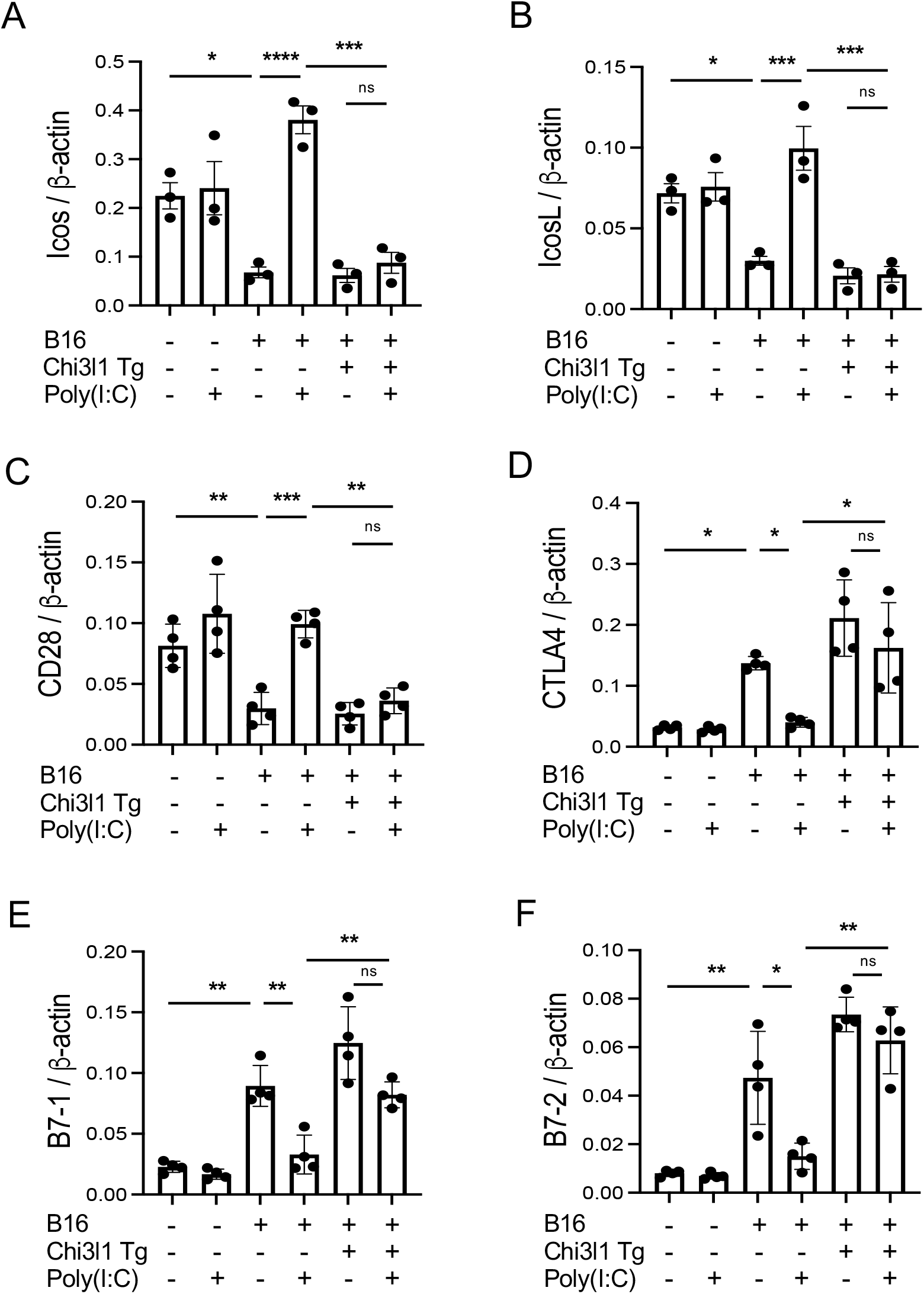
Effects of transgenic Chi3l1 on Rig-like helicase (RLH) inhibition of co-stimulation and induction of CTLA-4, B7-1 and B7-2. WT mice and Chi3l1 Tg+ mice were given B16-F10 (B16+) melanoma cells or PBS control (B16-) and treated with Poly(I:C) (Poly (I:C+) or its vehicle control (Poly(I:C-) and evaluated 2 weeks later. (A-F) RT-PCR was used to quantitate the levels of mRNA encoding ICOS, ICOSL, CD28, CTLA-4, B7-1 and B7-2 in lungs from WT mice and Chi3l1 Tg+ mice treated with Poly(I:C) (Poly (I:C+) or its vehicle control (Poly (I:C-) as noted. The plotted values in panels A-F represent the mean±SEM of the evaluations represented by the individual dots. \**P*<0.05, \*\**P*<0.01, \*\*\**P*<0.001. (ANOVA with multiple comparisons). ns, not significant.

### Antibodies against CHI3L1 and CTLA-4 interact to augment antitumor responses

Since CHI3L1 regulates co-stimulation and stimulates multiple ICP molecules including CTLA-4, we determined if anti-CHI3L1 and anti-CTLA-4 interact in antitumor responses. In these experiments, mice were treated with anti-CHI3L1 (FRG) and anti-CTLA-4 antibodies each alone or in combination. As shown in Figure 7, each anti-CHI3L1 and anti-CTLA-4 antibody inhibited melanoma lung metastasis when compared to isotype controls (Figure 7, A and B). Importantly, the antitumor responses in the mice treated with both antibodies in combination exceeded the effects in the mice treated with individual antibodies (Figure 7, A and B). These effects appeared to be at least additive in nature. These studies demonstrate that anti-CHI3L1 and anti-CTLA-4 interact to augment antitumor responses in lung melanoma metastasis.

**Figure 7.**
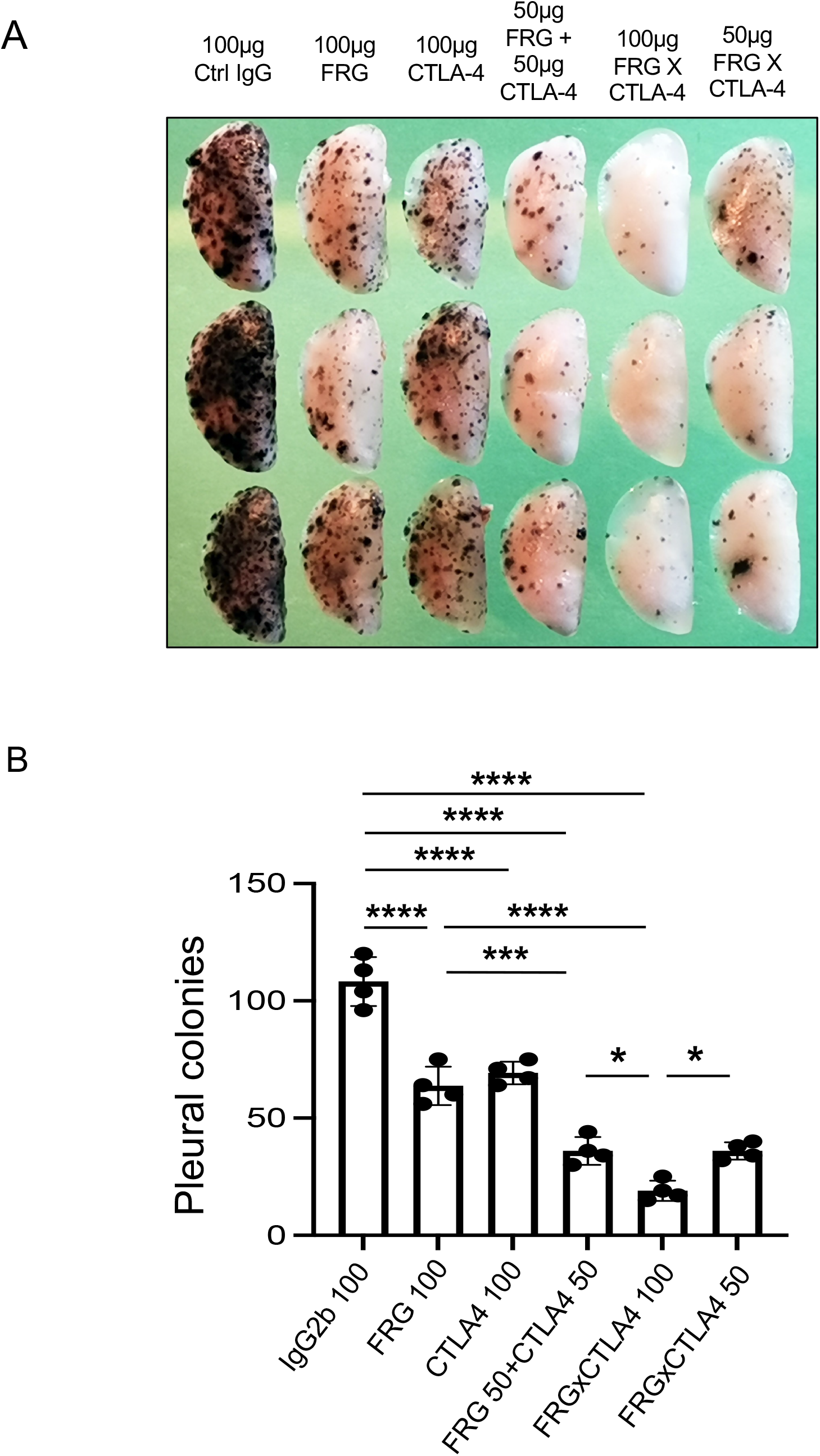
Anti-Chi3l1 and anti-CTLA-4 interact to augment antitumor responses in melanoma lung metastasis. WT mice were given B16-F10 (B16) melanoma cells or control vehicle and treated with control IgG, FRG and or anti-CTLA-4 antibodies, alone or in combination, and melanoma lung metastasis was evaluated 2 weeks later. (A) Representative lungs from mice treated with control IgG, FRG and or anti-CTLA-4 antibodies, alone or in combination. They were compared to the mice treated with indicated doses of the bispecific antibody FRG-CTLA-4. As noted, the antibodies were given at doses of 100μg or 50 μg every other day by intra-peritoneal injection. (B) The number of pleural melanoma colonies was quantitated in the lungs from the mice in panel A. Each dot is representative of an individual animal. Panel A is a representative of at least 3 evaluations. The values in panel B represent the mean±SEM of the evaluations represented by the individual dots in the lungs from the experiment representatively illustrated in panel A. \**P*<0.05. \*\*\**P*<0.001; \*\*\*\**P*<0.0001. (ANOVA with multiple comparisons).

### Bispecific antibodies that simultaneously target CHI3L1 and CTLA-4 synergistically induce CTL cell differentiation, PTEN expression and tumor cell death

In recent years it has become clear that combination therapy with ICP blockers can induce particularly potent responses in a variety of tumors including lung cancers (32–34). Recent studies have also demonstrated that bispecific antibodies can have powerful and or unique biologic effects compared to their individual component antibodies, alone and in combination (35). The studies noted above suggest that antibodies against CHI3L1 and CTLA-4 interact to enhance antitumor responses in melanoma metastasis. Thus, studies were undertaken to determine if bispecific antibodies that simultaneously target CHI3L1 and CTLA-4 (FRGxCTLA-4) initiate even more prominent antitumor responses. To answer this question, we generated bispecific antibodies by linking anti-CTLA-4 to anti-CHI3L1 (FRG) via its light chain (Figure S3). This bispecific FRGxCTLA-4 antibody manifest high and comparable levels of affinities to both CHI3L1 and CTLA-4 proteins (Kd=1.1×10^−9^) (Figure S3). Studies to undertaken to evaluate the effects of FRGxCTLA-4 to FRG and anti-CTLA-4 alone and in combination, in in vivo animal model of melanoma lung metastasis and in vitro cocultures composed of activated T cells and A357 human melanoma cells. In the in vivo experiments the bispecific antibodies were impressively powerful inhibitors of melanoma metastasis (Figure 7, A and B). These effects were synergistic in nature when compared to antibodies against CHI3L1 or CTLA-4 individually and more powerful antitumor effects were appreciated when the antibodies were administered simultaneously (Figure 7, A and B). In cocultures, each FRG and anti-CTLA-4 antibody significantly increased tumor cell apoptosis (Figure 8A). We noted additional increases in tumor cell death with simultaneous treatment of FRG and anti-CTLA-4 antibodies together, at least additive in nature (Figure 8 A). Importantly, we noted the highest levels of tumor cell apoptosis in the cells treated with the bispecific FRGxCTLA-4 antibody (Figure 8A). This effect of the bispecific antibody appeared to be synergistic in nature because the levels of tumor cell death were greatly exceeded the effects of individual FRG and anti-CTLA-4 antibody or in combination. In all cases, CD8+ cytotoxic T cells may play a significant role in this antitumor response because FRG and anti-CTLA-4 treatment enhanced T cell expression of CD8, perforin and granzyme expression and these effects were synergistically enhanced in cocultures with FRGxCTLA-4 treatment (Figure 8, B-D). Surprisingly, FRG and anti-CTLA-4, alone and in combination, also enhanced the expression of the tumor suppressor PTEN and these effects were also synergistically heightened in cocultures with FRGxCTLA-4 treatment (Figure 8E). In quantitative evaluation, the anti-tumor effects of mono and bispecific antibodies of FRG and CTLA-4 were all significant (Figure 8F). In sum, these studies highlight that bispecific antibodies that simultaneously target CHI3L1 and CTLA-4 have prominent additive or synergistic anti-tumor effects in vivo and in vitro via its ability to induce CD8+ cytotoxic T cells and enhance tumor cell expression of PTEN.

**Figure 8.**
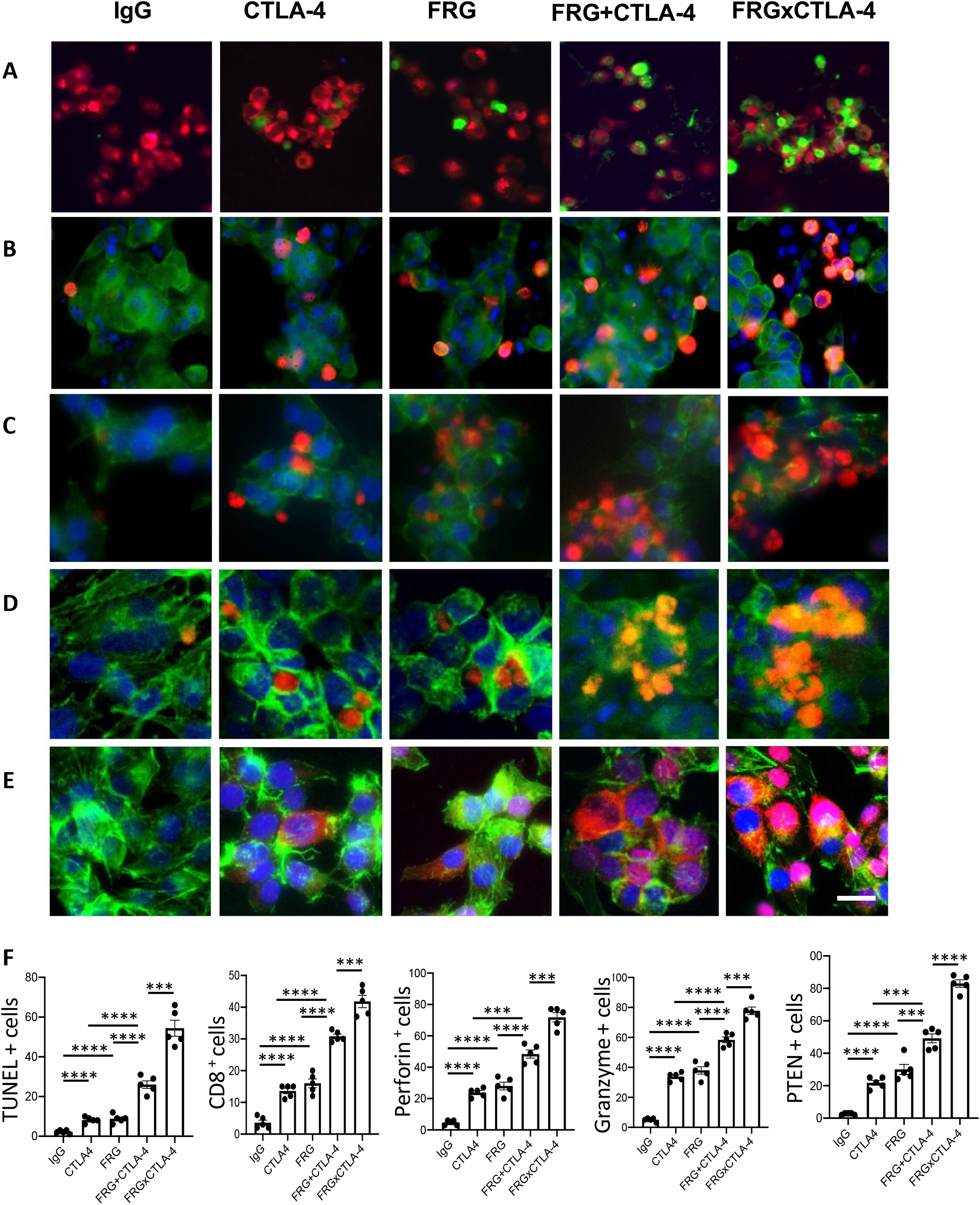
Bispecific antibodies that simultaneously target Chi3l1 and CTLA-4 induce synergistic CTL-mediated tumor cell death responses and tumor cell PTEN expression. The antitumor effects of the FRGxCTLA-4 bispecific antibody were evaluated in a co-culture system containing TALL-104 cells and A375 human melanoma cells. TALL-104 cells were activated by pretreatment with anti-CD3 and anti-CD28 (1 μg/ml each and incubated for 2 hrs in 5% CO2 and air at 37°C). The TALL-104 cells were then co-cultured with A357 human melanoma cells for 24 hours. These co-cultures were undertaken in the presence of the following antibodies; isotype control antibody (5μg/ml), anti-CTLA-4 or anti-Chi3l1 (FRG) alone (5μg/ml), or in combination (2.5 μg/ml each), and the bispecific FRGxCTLA-4 antibody (5μg/ml). (A) Representative demonstration and quantitation of apoptotic tumor cell death using in situ cell detection kit-fluorescein dUTP. TUNEL (+) cells stain green. (B-D) Representative demonstration and quantification of TALL-104 cell expression of CD8 (B), perforin (C) and granzyme (D). Tumor cells are green and positive staining TALL-104 cells are yellow-orange. (E) Representative demonstration and quantification of tumor cell PTEN. Tumor cells are green and PTEN is yelloworange. (F) Quantification of the evaluations in A-E. The % of TUNEL + tumor cells (Column A), % of TALL-104 cells expressing CD8 (column B), perforin (column C) and granzyme (column D) and % of tumor cells expressing PTEN (column E) are illustrated. These evaluations were done using fluorescent microscopy (x20 of original magnification). In these quantifications, 5 randomly selected fields were evaluated. The values in panel F are the mean ± SEM. \*\*\**P*<0.001, \*\*\*\**P*<0.0001. Scale bar =10μm, applies to all subpanels of A-E.

## Discussion

The immune surveillance hypothesis proposed in the 1950’s that the immune system could recognize and reject cancer cells as foreign (36). It is now known that CD8+ cytotoxic T cells are the main players driving adaptive immune responses to cancer (37) and that tumors get shielded from immune elimination by decreasing T cell co-stimulation and or expressing ligands that normally interact with inhibitory immune receptors that protect “self” (36). As a mechanism that mediates immunosuppressive responses in tumor or tumor microenvironment, we hypothesis that CHI3L1 plays a critical role in these immunosuppressive events mediated by immune checkpoint molecules. These studies revealed the ability of melanoma lung metastasis to inhibit T cell co-stimulation and induce CTLA-4 and its ligands. We further demonstrated that the effects of CHI3L1 are tumor independent, at least in part, because transgenic overexpression of CHI3L1, in the absence of tumor, reproduced these events. These findings suggest that immune checkpoint regulation is a potential mechanism of immunosuppressive effects of CHI3L1 that have been previously described in breast cancer (38). It is also reasonable to speculate that type I immune responses can be coopted to shut down antitumor immune responses potentially through this this mechanism (2, 39). They also provide significant therapeutic implications because simultaneous targeting of CHI3L1 and CTLA-4 using monoclonal antibodies in combination generate, at least additive, antitumor responses *in vivo* and induction of synergistic tumor cell death and CD8+ cytotoxic T lymphocyte differentiation with bispecific antibodies that simultaneously target CHI3L1 and CTLA-4.

The CD28 costimulatory receptor is a 44kDa membrane glycoprotein that is expressed on nearly all human T cells at birth (40). Binding of the CD28 receptor provides an essential second signal alongside TCR ligation which induces T cell activation (37). CD28 is the prototype of a family of co-stimulators that includes ICOS and others (37, 41). CD28 and CTLA-4 compete for the same ligands (B7-1 and B7-2). They regulate immune responses by providing opposite effects with CD28 stimulating and CTLA-4 inhibiting T cell function (37). In keeping with these findings, CD28 agonists have been developed that awaken T cells from a tolerant state and activate the immune system in the treatment of cancer and infection (42, 43). Our studies add to our understanding of CD28 and related moieties by demonstrating, for the first time, that CHI3L11 plays a critical role in their regulation. Specifically, they demonstrate that CHI3L1 inhibits ICOS, ICOSL and CD28 co-stimulation while augmenting the expression of the immune inhibitor CTLA-4. This supports the concept that combining checkpoint inhibitors and agonists of co-stimulation represents an exciting therapeutic approach for cancer therapy (44). It also demonstrates how anti-CHI3L1-based interventions can have these desired multifaceted effects.

Programed cell death (PD)-1 and CTLA-4 are co-inhibitory receptors that inhibit T cell activity and, as such, are referred to as immune checkpoints (45). Antibodies that block these receptors or their ligands have impressive clinical activity against an array of human cancers (45–47). In addition, the combination of anti-CTLA-4 and an anti-PD-1 or anti-CTLA-4 and an anti-PD-L1 are far more effective than either agent alone in clinical trials (42, 48) or pre-clinical models (49). In keeping with the importance of these interactions the mechanisms that underlie the interactions of CTLA-4 and the PD-1/PD-L1 axis have been studied. These recent investigations demonstrated that PD-L1 and CD80 (B7-1) bind to each other exclusively in *cis* and that this interaction blocks the immunosuppression induced by both the PD-1/PD-L1 axis and the CTLA-4/B-7 axis while leaving CD28/B-7 T cell co-stimulation unaltered (6). Our studies add to this mechanistic understanding by demonstrating that CHI3L1 stimulates both CTLA-4 and components of the PD-1/PD-L1 axis while simultaneously inhibiting ICOS and CD28 costimulation (6). When viewed in combination one can appreciate the multifaceted manner via which ICP moieties interact to suppress antitumor immune responses.

In tumors and the host microenvironment, activation of RIG-I/RLH signaling is a critical component of immune checkpoint inhibition (ICPI) induced by antibodies against CTLA-4 and PD-1, alone and in combination (30). The appropriate induction of ICPI triggers tumor cell death, presentation of tumor associated antigens by CD103+ dendritic cells and the accumulation of antigen specific CD8+ infiltrating T cells (30). It was speculated that the engulfed nucleic acids from disintegrating tumor cells in host myeloid cells can further enhance RLH pathway activation (30). In support of this notion, previously we demonstrated that RLH activation induced by Poly (I:C) stimulation powerfully inhibits the expression of CHI3L1 (29). The present studies also demonstrate that RLH inhibits CHI3L1, in turn, augments T cell co-stimulation and decreases the expression of CTLA-4. They also demonstrate that simultaneous treatment of anti-CHI3L1 and anti-CTLA-4 antibodies elicits, at least additive, antitumor effects and synergistic tumor cell death responses with the bispecific antibody treatment. They provide a potential mechanism of RIG-I/RLH activation that enhances the ICPI-mediated antitumor responses. Because CHI3L1 is a critical regulator of inhibitory immune checkpoint molecules and RIG-I/RLH activation inhibits CHI3L1, it is reasonable to speculate that CHI3L1 mediates the antitumor activities of RIG-I/RLH activation. If this is the case, the interventions that inhibit or block CHI3L1 can be used to maximize the therapeutic efficacy of immune checkpoint blockade.

Lung cancer is the leading cause of cancer deaths worldwide (50). In line with its importance, scientists developed new therapeutics based on tumor immunology for patients with primary and metastatic lung malignancies (36). In many cases these therapies are now first or second line lung cancer therapeutics (51). However, only a small number of patients responded to these therapies and the responses were often limited and not durable (52). To overcome these issues, various approaches including combination immunotherapy have been proposed. Among these, the immunotherapy with both anti-PD-1 and anti-CTLA-4 antibodies in combination has been most commonly employed and this combination demonstrated powerful effects in some malignancies (47). However, its effects in lung cancers has not been evaluated and potential toxicities can limit the use due to the exaggerated anti-self-immune responses induced by antibody combination (47). In keeping with the concept that combination immunotherapy may be therapeutically useful, we evaluated the effects of anti-CHI3L1 (FRG) and anti-CTLA-4 antibodies, alone and in combination. These studies demonstrate that the simultaneous treatment with anti-CHI3L1 and anti-CTLA-4 antibodies individually or bispecific antibodies that simultaneous target these two moieties result in additive or synergistic antitumor responses. Since RLH activation inhibits CHI3L1, we can also envision similarly augmented antitumor responses when RIG-I/RLH activators and anti-CTLA-4 antibodies are co-administered. Whether the therapeutic outcome and safety profile of these immunotherapeutic combinations are better or worse compared to anti-PD-1 plus anti-CTLA-4 antibody combination will need to be further determined.

Although antibody-based blockade of individual checkpoints has proven therapeutically useful in a variety of human cancers, the responses that are elicited are often not durable (53). This is in keeping with recent observations that demonstrate that the blockade of a single checkpoint target can lead to the compensatory upregulation of other checkpoint receptors (8, 54). As a result, many have turned to combinations of therapeutics that target non-redundant immune pathways to achieve enhanced antitumor responses (8). Our studies support this concept by demonstrating that the combined use of antibodies that target CHI3L1 (a multifaceted stimulator of ICP) and CTLA-4 results in enhanced antitumor responses. Additional studies will be required to further define the optimal combination of immune blockade antagonists for primary and metastatic lung cancer.

Bispecific antibodies are a growing class of immunotherapies with the potential to further improve therapeutic efficacy and safety (35). To determine if the bispecific antibodies that target CHI3L1 and CTLA-4 particularly effective, we generated bispecific antibodies and their antitumor effects in a T cell-melanoma cell coculture system were evaluated and compared to individual antibodies, alone or in combination. These studies demonstrate that bispecific FRGxCTLA-4 antibody synergistically induces tumor cell death compared to the effects of the individual antibodies. They also demonstrated a remarkable ability of FRGxCTLA-4 antibody to induce CD8+, perforin +, and granzyme+ cytotoxic T cells, that might cause the tumor cell death responses. These findings well aligned with a fundamental principle of tumor immunology, that host cytotoxic T cells can eliminate tumor cells (55). The synergistic responses induced by FRGxCTLA-4 antibody further support our notion that multiple immunoregulatory pathways are implicated in this response. They demonstrate that interventions that inhibit the production and or effector functions of CHI3L1 can diminish the ability of CHI3L1 to block tumor cell death and augment the expression of IFNα/β, ChemR23, phosphorylated cofilin and LimK2 (29). In previous studies, we also appreciated that CHI3L1-based interventions augment the expression of the tumor suppressor PTEN and type I immune responses and decrease M2 macrophage differentiation (29). All together, these studies demonstrate that bispecific antibodies that target CHI3L1 and CTLA-4 have impressive antitumor effects while inducing CD8+ cytotoxic T cell differentiation and tumor cytotoxicity.

In conclusion, these studies demonstrate that CHI3L1 inhibits the T cell co-stimulation induced by ICOS and CD28 and their ligands while stimulating the checkpoint inhibitor CTLA-4. They also demonstrate an additive antitumor response in melanoma lung metastasis with simultaneous treatment with individual antibodies against CHI3L1 and CTLA-4 and synergistic CD8+ cytotoxic T cell-mediated tumor cell death with bispecific antibodies that target these two moieties. These findings strongly support our notion that interventions that simultaneously target CHI3L1 can augment the therapeutic efficacy of immune checkpoint blockade of CTLA-4 in lung cancer and other malignancies. Additional studies of the biology and therapeutic outcomes of interactions between CHI3L1 and immune checkpoint inhibitors are warranted.

## Materials and methods

### Genetically modified mice

Mice with null mutations of CHI3L1 (*Chil1*^-/-^) and lung-specific CHI3L1 overexpressing transgenic mice (*Chi3l1 Tg*) have been generated and characterized by our laboratory as previously described (14, 56). All animals were humanely anesthetized with Ketamine/Xylazine (100mg/10mg/kg/mouse) before any intervention. The protocols that were used in these studies were evaluated and approved by the Institutional Animal Care and Use Committee (IACUC) at Brown University.

### Western blot analysis

20 to 40 μg of total lung lysates were prepared with RIPA lysis buffer (ThermoFisher Scientific, Waltham, MA, USA) and subjected to the evaluations according to the procedures that we recently reported (30). Anti-mouse ICOS (7E.17G9, Bio X Cell) and anti-mouse ICOSL (HK5.3, Bio X Cell), anti-mouse CD28 (37.51, Bio X Cell), anti-human-CTLA-4 (922101, R&D), anti-mouse-CTLA-4 (a generous gift from Prof. Lieping Chen, Medical Oncology, Yale School of Medicine), anti-mouse B7-1 (F-7, Santa Cruz Biotechnology) and anti-mouse B7-2 (BU63, Santa Cruz Biotechnology) and β-actin (C4, Santa Cruz Biotechnology) were used as primary antibodies and immunoreactive bands were captured and analyzed using a ChemiDoc (Bio-Rad) imaging system.

### RNA extraction and Real-time qPCR

Total lung mRNA was isolated using TRIzol reagent (ThermoFisher Scientific) followed by RNA purification using RNeasy Mini Kit (Qiagen, Germantown, MD) according to the vendor’s instructions. Semiquantitative real time (RT)-qPCR was used to measure the levesl of mRNA expression as described previously (13, 14). The sequences of primers used in these studies are presented in Table S1. Ct values of the target genes were normalized to the internal housekeeping gene β-actin.

### Generation of bispecific antibodies against CHI3L1 and CTLA-4

The anti-CHI3L1 monoclonal antibody (FRG) was generated as described previously (30). Bispecific CHI3L1 and CTLA-4 antibody (FRGxCTLA-4) was generated by a construct that links anti-CTLA-4 to the light chain of Fc portion of FRG antibody (Figure S3). HEK-293T cells were transfected with the bivalent FRGxCTLA-4 construct using Lipofectamine^™^ (Invitrogen, CA, USA). The bispecific antibody was purified from the collected supernatant using a protein A column (Thermo Scientific # 89960; IL, USA) and ELISA was used to assess ligand binding affinity and sensitivity of the antibody. The original murine anti-CTLA-4 monoclonal antibody (57) was a generous gift from Prof. Lieping Chen (Medical Oncology, Yale School of Medicine). Limulus Amebocyte Assay (Pierce, Rockford IL) was used to exclude endotoxin contamination.

### Melanoma lung metastasis and antibody treatment

B16-F10 (B16) murine melanoma cells were purchased from ATCC (Cat#: CRL-6475, Manassas, VA) and maintained in Dulbecco’s Modified Eagles Medium (DMEM) supplemented with 10% Fetal bovine Serum (FBS) and 1% penicillin & streptomycin as recommended. Anti-CHI3L1 (FRG), anti-CTLA-4, bispecific antibodies (FRGxCTLA-4) and its isotype control IgG (IgG2b) were delivered via intraperitoneal injection, alone or in combination, every other day from the day of the B16 tumor cell challenge for 2 weeks (28, 29). Melanoma lung metastasis was quantified by counting the melanoma colonies on the pleural surface as described previously (28, 29).

### Fluorescence-activated cell sorting (FACS) analysis

Whole mouse lungs were digested and single cell suspension was prepared using dissociation kit (Miltenyi Biotec, Auburn, CA) as per the vendor’s instructions. Cells were labeled with fluorescent antibodies against CD45 (30-F11, Fisher Scientific), CC10 (E-11, Santa Cruz Biotechnology), SPC (H-8, Santa Cruz Biotechnology), CD3-APC (145-2C1, Fisher Scientific), CD11b-FITC (M1/70, Fisher Scientific), MHC-II (M5/114.15.2, Fisher Scientific), ICOS-PE-cy7 (C398.4A, Biolegend), CD28-PE-cy7 (25-0289-42, ebioscience), and CTLA-4-PE-cy7 (25-1522-82, ebioscience). BD FACSAria IIIu and FlowJo V10 software were used for flowcytometry data collection and analysis, respectively. The gating strategy of CD3(+) and CTLA(+) Immune cells is illustrated in Figure S4.

### Immunohistochemistry

Formalin-fixed paraffin embedded (FFPE) lung tissue blocks were serially sectioned at 5μm-thickness and were subjected to immunohistochemistry (IHC) evaluations according to the procedures that we recently reported (30). The primary antibodies used in this IHC evaluations are: α-CTLA-4 (obtained from Prof. Lieping Chen, Yale University), α-CC10 (E-11, Santa Cruz Biotechnology), α-SPC (M-20, Santa Cruz Biotechnology), and α-CD68 (#ab125212, Abcam).

### Melanoma cells and TALL-104 T cell co-cultures

Human melanoma A375 cell line (CRL-1619) and TALL-104 T cells (CRL-11386) were purchased from ATCC and maintained according to the protocols provided by the vendor. The TALL-104 cells were first stimulated with anti-human CD3 antibody (5μg/ml) (Biolegend) and anti-human CD28 antibody (5μg/ml) (#MABF408, Millipore). Then, the activated TALL-104 cells and A375 melanoma cells were resuspended together at a 6:1 ratio in complete RPMI media, dispensed into the multi-well slide chambers and incubated for 1 hour in 5% CO2 at 37°C then isotype control or testing antibodies were added to the co-cultured cells. The effects of the isotype control antibodies were compared to the effects of antibodies against CHI3L1 (FRG; 5μg), CTLA-4 (5μg), FRG+CTLA-4 administered simultaneously (2.5μg each) and FRGxCTLA-4 (5μg). After 48 hours of incubation, cells were subjected to TUNEL and immunocytochemistry evaluations using fluorescent-labeled antibodies against CD8, perforin, granzyme-B and PTEN by as previously described (30).

### Quantification and Statistical analysis

Statistical significance and differences between two groups were compared with 2-tailed Student’s *t* test and one-way ANOVA with Turkey post hoc test was used for multiple group comparison. A *P* value less than 0.05 was considered statistically significant.

## Author Contributions

Conception and design: BM, CGL, JAE, CGL; Generation of experimental resources and data collection: BM, SK, BA, HK, CML; Analysis and interpretation: BM, SK, CGL, JAE; Drafting the manuscript for important intellectual content: BM, CGL, JAE

## Acknowledgements

This work was supported by National Institute of Health (NIH) grants PO1 HL114501(JAE), and R01 HL155558 (CGL) from NHLBI.

## Competing Interests

JAE is a cofounder of Elkurt Therapeutics and is a stockholder of, and serves on the Scientific Advisory Board for Ocean Biopharma, Inc., which develops inhibitors of 18 glycosyl hydrolases as therapeutics. JAE, CGL and SK have composition of matter and use patents relating to antibodies against CHI3L1. The other authors have declared that no conflict of interest exists.

## Notes

### Competing Interest Statement

Dr. Jack A Elias is a cofounder of Elkurt Therapeutics and is a stockholder of, and serves on the Scientific Advisory Board for Ocean Biopharma, Inc., which develops inhibitors of 18 glycosyl hydrolases as therapeutics. Drs. Jack A Elias, Chun Geun Lee and Suchitra Kamle have composition of matter and use patents relating to antibodies against CHI3L1. The other authors have declared that no conflict of interest exists.

### Summary of Updates

Overall manuscript has been substantially revised in the main text, figures, and supplemental data and figures in the process of revision after submission to Frontiers in Immunology (In Press)

