## Supplemental data and figures for "CHI3L1 enhances melanoma lung metastasis via regulation of t cell co-stimulators and CTLA-4/B7 axis"

### Supplementary Table

Table S1. Sequences of RT-PCR primers used in this study

| Gene | Sequence (5'to 3') | Length |
|---|---|---|
| Icos-S | atg aag ccg tac ttc tgc cg | 20 |
| Icos-AS | cgc att ttt aac tgc tgg aca g | 22 |
| IcosL-S | taa agt gtc cct gtt ttg tgt cc | 23 |
| IcosL-AS | att gca ccg act tca gtc tct | 21 |
| CD28-S | gtt ctt ggc tct caa ctt ctt ct | 23 |
| CD28-AS | tga ggc tga cct cgt tgc tat | 21 |
| CTLA4-S | ttt tgt agc cct gct cac tct | 21 |
| CTLA4-AS | ctg aag gtt ggg tca cct gta | 21 |
| B7-1-S | acc ccc aac ata act gag tct | 21 |
| B7-1-AS | ttc caa cca aga gaa gcg agg | 21 |
| B7-2-S | ctg gac tct acg act tca caa tg | 23 |
| B7-2-AS | agt tgg cga tca ctg aca gtt | 21 |
| Actin-S | aga ggg aaa tcg tgc gtg ac | 20 |
| Actin-AS | caa tag tga tga cct ggc cgt | 21 |

### Supplemental Figures

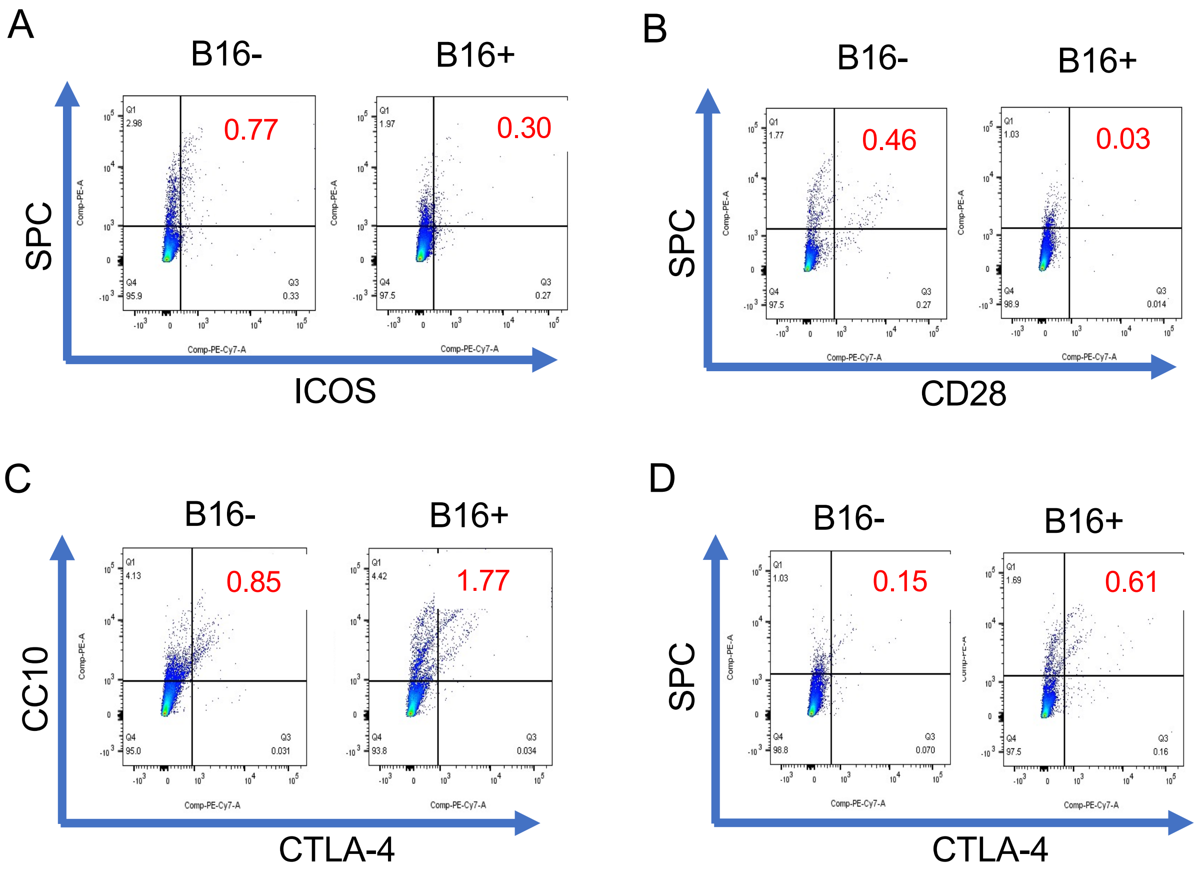


**Figure S1. Expression of ICOS, CD28, CTLA4 in lung epithelial cells in melanoma lung metastasis.** FACS evaluations of ICOS, CD28, and CTLA-4 expression using airway epithelial (CC10) and alveolar (SPC) epithelial cell specific markers in the lungs with melanoma (B16+) compared to vehicle (B16-) challenged mice. This is representative of a minimum of 2 similar evaluations.

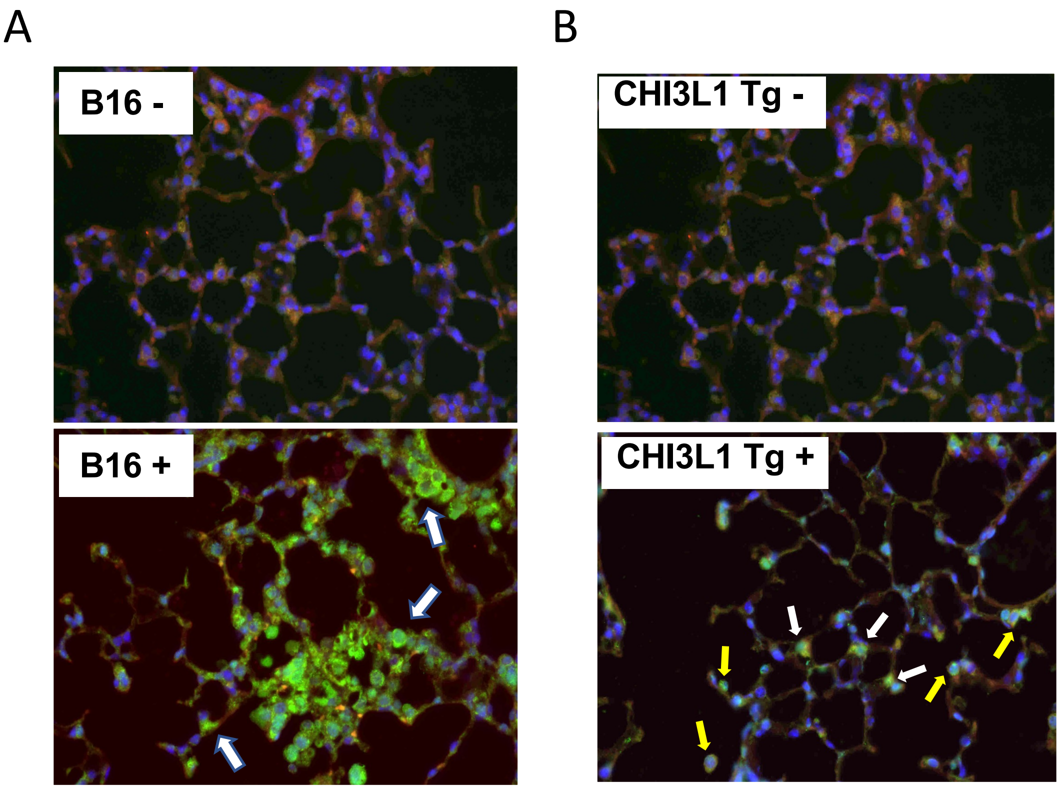


**Fig. S2. CTLA-4 expression in melanoma lung metastasis and in the lungs of CHI3L1 overexpressing transgenic mice**. (A) Increased expression of CTLA-4 (green, arrows) in the lungs challenged with B16 melanoma cells (B16+) compared to non-challenged ones (B16-). B) Increased expression of CTLA-4 (green) in the lungs of Chi3l1 transgenic mice (Tg+) compared to wild type (Tg-) lungs. Based on the location of the cells and morphology, the alveolar epithelial cells and macrophages/monocytes with CTLA (+) staining were identified and indicated by white and yellow arrows, respectively. Representative fluorescent immunohistochemical staining, x40 magnification.

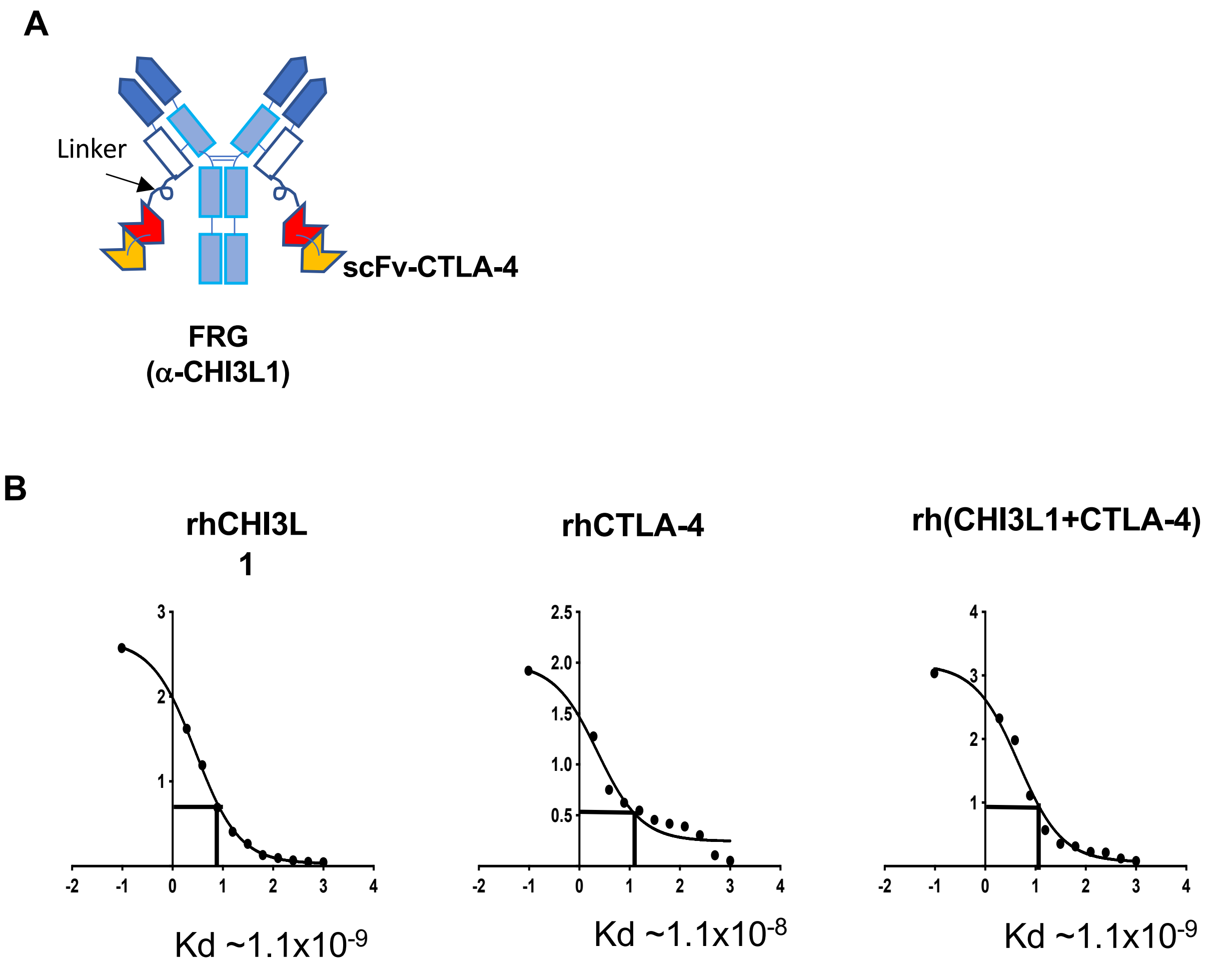


**Figure S3. The structure and the binding affinity of FRGxCTLA-4 bispecific antibody**. (A) Schematic illustration of the structure of the bispecific antibody FRGxCTLA4 in which anti-CTLA-4 is linked to FRG via its light chain. (B) The affinity of FRGxCTLA-4 antibody was evaluated by competitive ELISA against recombinant human (rh) Chi3l1, rhCTLA4 and mixture of rhCHI3L1 and rhCTLA-4.

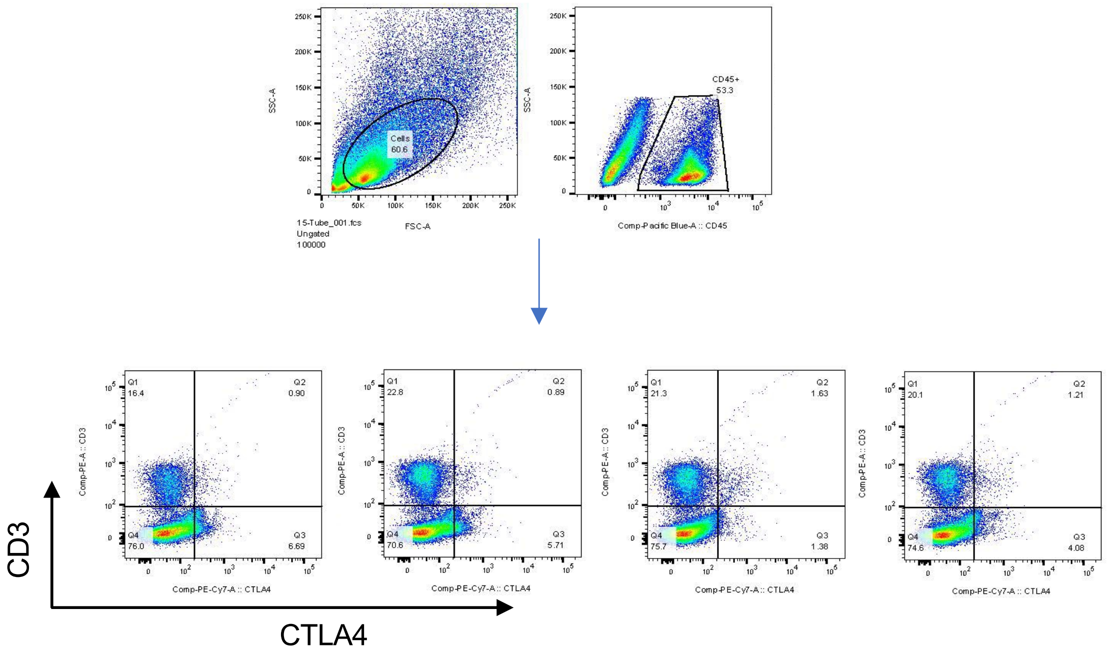


**Fig. S4. Gating strategy of CD3(+)/CTLA4(+) cells in the lung**. The live cells and CD45(+) cells from the lungs of individual mice were gated first then the CD3(+)/CTLA4(+) cells were counted and plotted.
